# Systemic organellar genome reconfiguration along the parasitic continuum in the Broomrape family (Orobanchaceae)

**DOI:** 10.1101/2025.06.03.657626

**Authors:** Yanlei Feng, Susann Wicke

## Abstract

The transition from autotrophy to heterotrophy in parasitic plants disrupts organellar coordination and presents a unique opportunity to examine the coevolution of cellular genomes. Using the Broomrape family (Orobanchaceae) as a model, we analyzed mitochondrial and plastid genome evolution across 30 species representing the full spectrum of parasitic lifestyles. We show that plastid genome reduction is correlated with mitogenomic expansion, revealing a striking inverse relationship between genome compaction and inflation. Mitogenome enlargement in parasitic taxa is driven by the accumulation of horizontally and intracellularly transferred DNA, proliferation of short repeats, and integration of unique sequences with no detectable homology. Notably, foreign DNA insertions converge toward native mitochondrial GC content and often retain structural functionality, suggesting selective maintenance. Relaxed selection in ATP synthase and ribosomal genes contrasts with intensified selection on components of electron transport and cytochrome c maturation, reflecting functional reconfiguration of mitochondrial respiration in parasitic plants. RNA editing, intron loss, and frameshift insertions further reshape gene structure, particularly in obligate parasites. Together, our findings suggest that parasitism initiates a systemic genomic feedback loop in which relaxed selection and disrupted maintenance mechanisms affect even distant genomic compartments. This study provides the first comprehensive evolutionary framework for multi-compartment genome remodeling in parasitic plants and highlights the dynamic interplay between lifestyle specialization and organelle genome evolution.

## Introduction

Plants maintain two semi-autonomous genetic systems to control the function and crosstalk between plastids, mitochondria, and the nucleus. Each of the organellar genomes encodes incomplete sets of proteins for their respective primary functions in addition to several housekeeping elements. The remainder of proteins needed for photosynthesis, cell respiration, or the housekeeping machineries are under the control of the nuclear genome and imported back into the organelles. The intricate nature of multi-subunit protein complexes requires its subunits to evolve in concert to compensate for potential deleterious mutations. However, it remains unknown what and how fast coevolutionary responses come about upon a systemic disturbance within and between functional complexes, let alone across cellular genomes.

Parasitic plants are an excellent model system to elicit and pyramid the molecular (co)evolutionary interdependence within and between genome compartments and lifestyle changes. The ability to live of a heterotrophic carbon source has evolved many times independently in land plants (reviewed in Wicke and Naumann 2018; Nickrent 2020). While the so-called mycoheterotrophic plants epiparasitize mycorrhizal networks, haustorial parasites develop a specialized feeding organ, the haustorium, to build a direct physiological bridge to another plant. Water, nutrients, and macromolecules including proteins and nucleic acids channel through the haustoria (Haupt et al. 2001; Aly et al. 2011; LeBlanc et al. 2012) to either supplement the reservoirs of a photosynthetic parasite temporarily (facultative hemiparasite) and during critical life stages (obligate hemiparasite), or to completely provide all nutrition when there is no own photosynthetic activity (holoparasites) (lifestyle definitions *sensu* Wicke and Naumann 2018). Parasitism relaxes the selectional constraints on photosynthesis genes, resulting in rampant gene loss, a gradual acceleration of substitution and indel rates in retained genes, as well as considerable micro- and macrostructural reconfigurations in plastid genomes (plastomes) (e.g., dePamphilis and Palmer 1990; Wicke et al. 2013; Feng et al. 2016; Barrett et al. 2018).

Genomic changes in parasitic plants extend beyond plastomes, but their often inaccessibly large genomes (e.g., Wicke 2013; Yoshida et al. 2019; Lyko and Wicke 2021) continue to hamper evolutionary-genomic surveys. In line with the massive morphological changes in parasitic plants, the transition to a nonphotosynthetic way leaves footprints also on many other molecular pathways. This assumption is corroborated by the complex patterns of gene family diversification and reduction in some nuclear genomes of heterotrophic plants (Sun et al. 2018; Vogel et al. 2018; Yoshida et al. 2019; Cai et al. 2021; Xu et al. 2022).

The shift away from photo-autotrophic energy conversion and the specialization on a heterotrophic energy source likely affects the mitochondrial respiration complexes notably. Mitochondrial respiration supplies energy in plants when photosynthesis does not, so we would expect a high selection pressure to act on respiration-related genes, a critical proportion of those encoded in plant mitochondrial genomes. However, rather than an intensification of selectional constraints in respiratory complexes, we have observed functional losses in mitochondrial genomes (mitogenomes) of some parasitic plants from independent lineages (Molina et al. 2014; Skippington et al. 2015, 2017; Fan et al. 2016), including the complete loss of complex I of obligate parasitic mistletoes (Maclean et al. 2018; Senkler et al. 2018). In contrast, rather large mitogenomes were seen holoparasitic *Cynomorium coccineum* (1.11 Mbp, Cynomoriaceae, Bellot et al. 2016), *Lophophytum mirabile* (822 Kbp, Balanophoraceae, Sanchez-Puerta et al. 2017), *Cuscuta japonica* (814 Kbp, Convolvulaceae Lin et al. 2022), and *Cistanche* (>1,700 Kbp, Orobanchaceae; Miao et al. 2022). But since, in contrast to plastomes, mitogenomes evolve highly dynamically in plants in general (e.g., Mower 2020), it remains to be established whether the reported mitogenomic and functional reconfigurations are linked to parasitism *per se*. Similarly, although the intimate connection between parasitic plants and their hosts likely facilitates the incorporation of foreign gene fragments into parasite mitogenomes (e.g., Davis and Wurdack 2004; Mower et al. 2004, 2010; Barkman et al. 2007; Davis and Xi 2015), the many reports of ordinary (i.e., nonparasitic) plants containing mitochondrial DNA with genes from other species (rewiewed in Richardson and Palmer 2007; Hao et al. 2010; Rice et al. 2013), the lifestyle-associated higher susceptibility for horizontal or intracellular DNA transfer (HGT and IGT, respectively) remains to be shown in an evolutionary framework.

Here, we assess the series, relative timing, and interdependency of organelle genome reconfiguration during the transition from an autotrophic to a completely parasitic lifestyle in plants. Specifically, we test whether the transition to parasitism triggers interdependent of changes in the functional capacities, structures, and rates of molecular evolution of organelle genomes across genomic compartments, and we infer the evolutionary history of mitogenomic mosaics in parasitic plants. To this end, we reconstructed the plastid and mitochondrial genomes from 30 nonparasitic and parasitic species across the spectrum of trophic specializations of the Broomrape family (Orobanchaceae). Our analysis reveals that mitogenome evolution proceeds apparently reversely proportional to plastome reduction along the transition to holoparasitism. We can show that a lifestyle-dependent mitogenomic inflation in parasitic Orobanchaceae is due to increasing proportions of unique (“novel”) DNA and the partly rampant incorporation of foreign DNAs from other cellular genomes and potentially from host plants. Based on our findings, we suggest that the transition to obligate parasitism initiates a molecular evolutionary feedback loop, in which the relaxation of selection in one subgenome indirectly affects also other, even distant genomic compartments through shared repair and maintenance mechanisms, even though these as well as the DNA regions they maintain continue to evolve under purifying selection.

## Results

### Structural evolution of mitochondrial genomes in Orobanchaceae

Mitogenomes of Orobanchaceae vary extremely in size and structure (Fig. 1A and Supplementary Table S1). The nonparasitic *Lindenbergia philippensis* (344 Kbp) and the facultative parasite *Melampyrum pratense* (316 Kbp) are at the lower bound of mitogenome size, whereas the obligate parasites *Schwalbea americana*, *Striga gesnerioides*, *St. hermonthica* represent the upper bound with 2.67, 3.40, and 3.57 Mbp, respectively. Holoparasites belonging to the Broomrape clade (i.e., *Aphyllon*, *Conopholis*, *Kopsiopsis*, *Phelipanche*, and *Orobanche*) tend to have slightly bigger genomes, often with more than 900 Kbp, than their hemiparasitic relatives. We achieved the reconstruction of single mitogenome mastercircles for *L. philippensis* as well as for the facultative hemiparasites *Triphysaria versicolor*, like that in *Castilleja paramensis* (Fan et al. 2016). All other mitogenome assemblies resulted in the reconstruction of multiple mitochondrial fragments, most of which minicircles, with the holoparasite *Kopsiopsis hookeri* standing out with 20 circular molecules. However, hampered by high proportions of repetitive DNA, some mitogenome fragments in *Melampyrum pratense*, *Orobanche gracilis*, *Or. rapum-genistae*, *Schwalbea* , *St. gesnerioides*, and *St. hermonthica* appeared to be in a linear conformation. Identification of locally collinear blocks revealed short blocks of conserved mitochondrial DNA, whose locations to one another are highly variable across Orobanchaceae and even between close relatives, suggesting an exceedingly rapid evolution of mitogenome structure (Fig. 1B). In contrast, as parasitism unfolds, plastid genomes reduce in size and function and retain an overall stable chromosomal configuration with low numbers of repeats and rare events of structural rearrangements (Fig. 1A).

**Figure 1.**
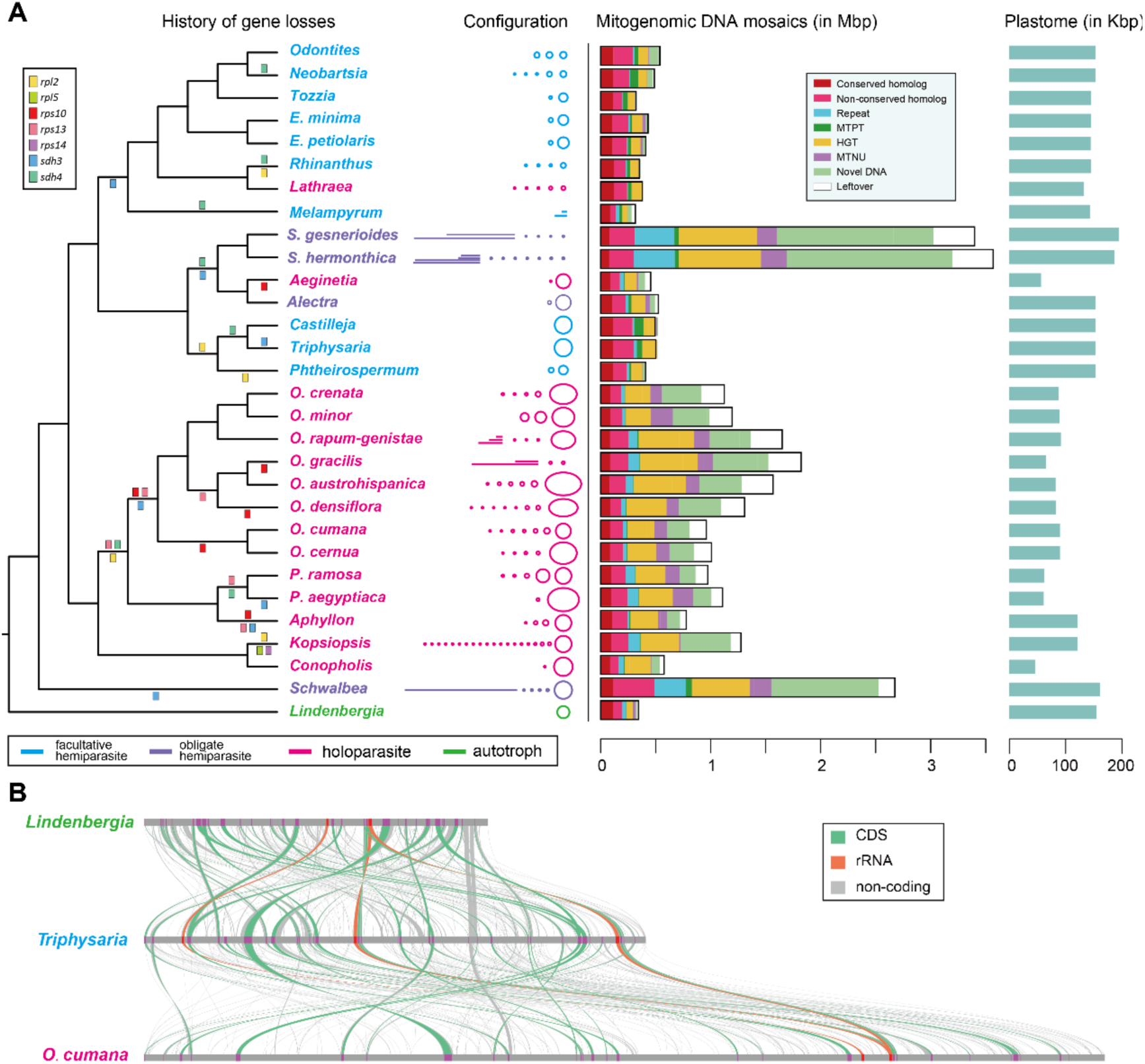
Gene loss, structural variation, and genome architecture of mitochondrial genomes in Orobanchaceae. **(A)** Gene loss events across the phylogeny. Colored blocks above and below branches indicate gene pseudogenization and deletion, respectively. Central schematics depict genome configurations and total lengths; circles and lines represent circular and linear mitochondrial chromosomes. Colored barcodes indicate putative genome components inferred from comparative analyses. Hatched areas within the “Novel” category denote regions shared among congeneric species. **(B)** Extensive mitochondrial genome rearrangements illustrated for three representative Orobanchaceae species.

We tested for associations between lifestyle changes and genomic reconfigurations of organelle DNA by comparing random-walk models under correlated and uncorrelated trait evolution using maximum likelihood. This analysis revealed that physical mitogenome inflation and plastid genome reduction is correlated significantly with the transition to parasitism and the nonphotosynthetic lifestyle (Likelihood ratio test, LRT: p-value <0.002). Longer branches contribute significantly to the evolution of these traits (LRT: p <0.001), whose rate of change weakly accelerates as time progresses (LRT: p <0.001). Notably, lifestyle and mitogenome size are positively correlated, whereas, in contrast, we correlations are negative for lifestyle and plastome reduction as well as between the sizes of the two organellar genomes. Moreover, considering also nuclear genome sizes in the discovered lifestyle-genome web, we also observe significant associations between all these traits (LRT: p-value <0.001), with gradual increases of nuclear and mitochondrial genomes compared to the reductive evolution of plastomes along parasitic specialization. That means, parasitic specialization progressing towards holoparasitism in Orobanchaceae coincides with a more dramatic reduction of plastomes, nuclear and mitogenomes both gain size in a gradual, non-adaptive mode of evolution.

### Changes of mitochondrial coding capacity in parasites

Orobanchaceae mitogenomes contain between 31 and 36 protein-coding genes (repeat genes excluded), 3 ribosomal RNAs (rRNAs) and 15-20 transfer RNAs (tRNAs) (Supplementary Table S1). 24 core mitochondrial genes (*atp1*, *4*, *6*, *8* and *9*, *ccmB*, *C*, *Fc* and *Fn*, *cob*, *cox1*-*3*, *matR*, *mttB*, *nad1*-*7*, *9,* and *4L*) and the ribosomal protein subunit genes *rpl10*, *rpl16*, *rps3*, *rps4* and *rps12* are present in all species (Fig. 2A). In hemiparasites, only *rpl2* is functionally or physically lost in a few taxa, whereas *rpl2*, *rps10*, and *rps13* have been lost in most holoparasitic species. *Kopsiopsis* is the only species that lost *rpl5* and *rps14*. Succinate dehydrogenase subunit genes *sdh3* and *sdh4* experienced independent losses in the parasites, but *Lindenbergia* and *Mimulus* both retain intact copies. This finding of independent losses indicates a potential functional transfer in the last common ancestor of parasitic Orobanchaceae, or earlier. In sum, despite their larger mitogenomes, holoparasites and a few hemiparasites encode fewer intact mitogenes.

**Figure 2.**
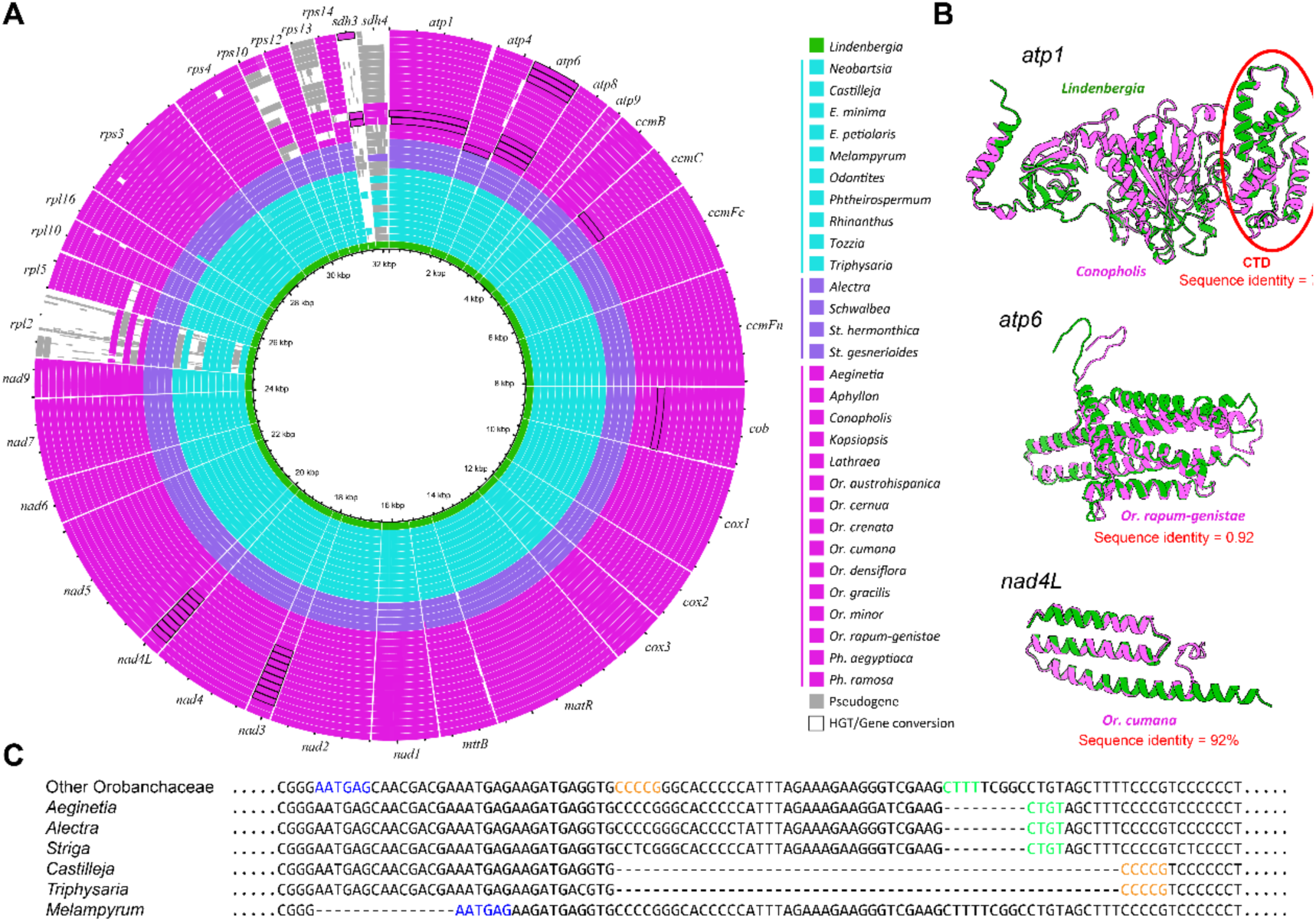
Mitochondrial gene variation in Orobanchaceae. **(A)** Mitochondrial protein-coding genes identified across Orobanchaceae species. Horizontally transferred ribosomal protein genes (*rps2*, *rps7*, *rps19*) are not shown. **(B)** Horizontally transferred genes (magenta) did not alter predicted protein structures. In *Conopholis*, the 3′ region of *atp1* shows evidence of gene conversion from Ericaceae (Supplementary Table S2), resulting in 77% nucleotide identity. Nevertheless, the predicted C-terminal domain (CTD) remains highly similar to that of *Lindenbergia* (green). Comparable patterns were observed for *atp6* in *Orobanche rapum-genistae* and *nad4L* in *Orobanche*. **(C)** Example of microhomology-mediated deletion in the *ccmFc* gene. Colored segments indicate putative microhomologous repeat pairs contributing to structural rearrangements.

Protein-coding genes in *Lindenbergia* and hemiparasites mitogenomes are rather conserved. In holoparasites, however, many genes reside have either an intact or pseudogenized extra copy and/or some of the parasites’ genes share a higher similarity with distantly related species than with Orobanchaceae. Blast searches as well as phylogenetic analyses indicate that several of otherwise lost genes (e.g., *sdh3*, *rps2*, *rps7*, *rps19*) have been re-gained by integration of a foreign fragment, likely originating from potential host plants (Supplementary Table S2). A sequence homology-based analysis suggest that all fragments were incorporated through DNA-mediated transfers. We also observe gene conversions between horizontally acquired and native mitogenes, resulting in mosaic open reading frames (Supplementary Table S2). Though the foreign and local genes exhibit divergence in the sequences, they demonstrate remarkable similarity in protein structure (e.g., Fig. 2B), which implies that HGT has not compromised the genes’ functionality.

Some protein-coding and ribosomal RNA genes also exhibit conspicuous insertions of deletions (indels). For example, a 55 bp insertion in *Melampyrum rrnS* is most similar to an early-diverging eudicot, and many particularly short repeat fragments occur in genes of *Aeginetia* and *Striga* (see below). In contrast, *ccmFc* of *Castilleja* and *Triphysaria* both have a 54 bp deletion, and *Or. rapum-genistae* has 87 and 120 bp deletions in *rps3* and *rps4*, respectively. We found microhomologous sequences to reside at such deletion sites, implying that those have originated through microhomology-mediated end joining (MMEJ) (Fig. 2C). It is noteworthy to mention that (functional) gene incorporations and sequence reconfigurations more commonly occur in obligate parasite and especially in holoparasites, with the exception of *Lathraea*, which represents the youngest completed transition to a nonphotosynthetic lifestyle in Orobanchaceae.

The intron number within genes are similar to other Lamiales relatives: *Trans*-splicing introns exist in *nad1*, *nad2*, and *nad5*; *ccmFc*, *cox1*, *cox2*, *rps3*, and *rps10* have one *cis*-splicing introns, whereas *nad4* and *nad7* both harbor three. Where present, the *rpl2* intron has been lost in all the species, and the intron *cox2*i700 has been lost from nine species in two independent events (Supplementary Table S1). The only group I intron in *cox1* exists in all Orobanchaceae species, including the autotrophic *Lindenbergia* and *Rehmannia* (Han et al. 2024).

### Expansion of unique DNA and incorporation of foreign DNA fragments

Homologous DNA in Orobanchaceae mitogenomes (i.e., CH and NCH) accounts for 80 and 200 Kbp, of which ∼55 Kbp are found in all taxa by all (Supplementary Table S3). These regions correspond mostly to genic regions as well as three intergenic but expressed spacers close to *trans*-spliced exons. The lengths of the homologous DNA is uncorrelated with the overall mitogenome lengths (PGLS P-value = 0.456).

HGT-like DNA incorporations correlate significantly with mitogenome size in Orobanchaceae (PGLS, P-value = 0.001; Figs 1A and 3A), suggesting that incorporation of foreign DNAs contribute to mitogenomic inflation. Our analyses show that Orobanchaceae mitogenomes consist of complex DNA mosaics of rampantly incorporated DNA (Supplementary Table S3). Between 10 and 32% of the mitogenomes originate from plastid or nuclear fragments or DNA from other, potential host plants. Of the latter, mitochondrial DNA of other species account for up to 95% of the length of non-native DNA in Orobanchaceae (Supplementary Tables S3). Foreign DNA from other plants mostly correspond to the parasites’ host preferences (Fig. 3B Supplementary Tables S4). For instance, *Aeginetia* contains considerable large amounts of DNA with highest similarity to Poaceae, whereas many *Orobanche* specie harbor Fabaceae-like fragments. However, some apparently non-native DNA also originate from less common host families, like Malvaceae in *Orobanche*.

**Figure 3.**
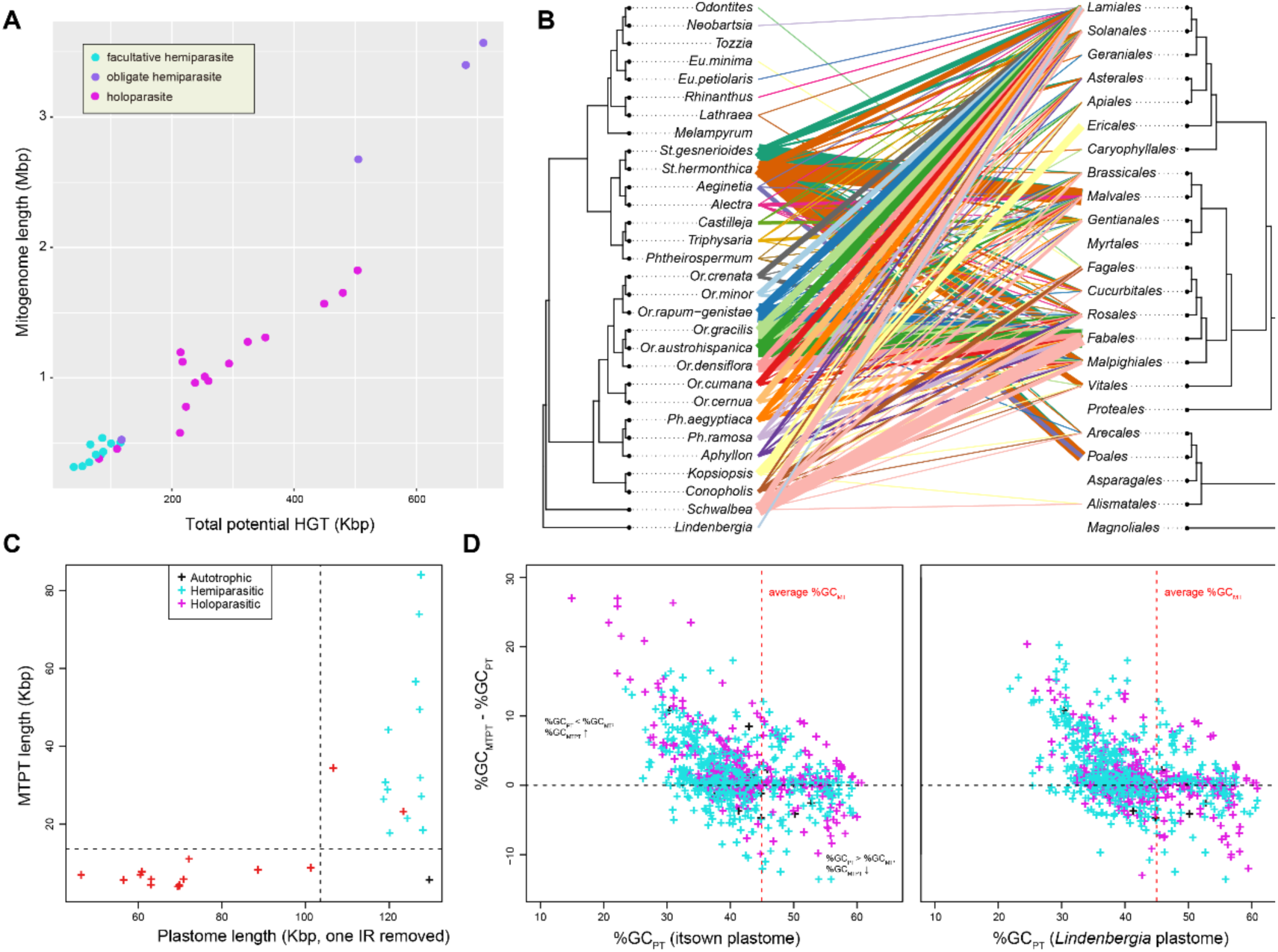
Distinct patterns of horizontal gene transfer (HGT) and mitochondrial plastid DNA transfer (MTPT) in holoparasitic Orobanchaceae. **(A)** Total HGT length is positively correlated with mitogenome size. **(B)** HGT size and inferred donor lineages. Putative donor species were grouped by order (right); connecting lines indicate likely donors, with line thickness proportional to transferred sequence length (>10 kbp only). Each recipient species is color-coded. **(C)** MTPTs (mitochondrial plastid DNA transfers) are longer in species with larger plastomes (only one inverted repeat counted). (B′, C′) GC content (%GC) of plastid-derived mitochondrial sequences shows convergence after transfer. (B′) Comparison with native plastome of the same species; (C′) comparison with the *Lindenbergia* plastome.

To disentangle the direction of intracellular DNA transfer, we used a combined database of multiple mitochondrial and nuclear genomes plus subsequent phylogenetic inference. Even though not all transfer directions could be resolved unambiguously, our analysis showed that holoparasites contain more DNA from their nuclear genome than hemiparasites (Fig. 1A, Supplementary Table S3 and S5). Mitochondrial insertions of nuclear DNA are normally rather short (∼1 Kbp) and the similarity between the corresponding mitogenomic and nuclear pairs is around 80%, although also identical pairs exist. We also found that hemiparasites and *Lindenbergia* contain considerably longer sequences from mitochondrial DNA in their nuclear genome (NUMTs), perhaps indicating a stronger interaction between these two genomic compartments.

After incorporation of intracellular transferred DNA, we observe a convergence of nucleotide composition. The lengths of inserted plastid-derived sequences are associated mitogenome size (Supplementary Table S3), but we noticed that species with longer plastid genomes contain more and longer plastid inserts in their mitogenomes (Fig. 3C). For example, plastid-derived DNA in hemiparasites is usually long and have a higher identity to their plastid counterparts, potentially indicating more recent or ongoing transfers. The plastid-derived fragments of Orobanchaceae mitogenomes diverge from true plastid DNA regarding their overall sequence (e.g., single nucleotide polymorphisms (SNPs); microstructural changes) as well as their nucleotide composition. Plastome-specific GC contents of Orobanchaceae are normally less than 38.5 %, compared with 43.5–45.6 % in mitogenomes (Supplementary Table S1). Plastid-derived mitochondrial DNAs, however, show a significant convergence to mitochondrial GC levels (PGLS, P-value = 0.001; Fig. 3D), even when accounting for the reduction GC contents in holoparasites by using the *Lindenbergia* plastome as null control for an undegraded reference for all taxa (PGLS, P-value = 0.001; Fig. 5c). The convergence trend is bidirectional, meaning that the GC content of plastid DNA with originally more AT bases increases, fragments with higher-than mitogenomic GC occurrences (e.g, *rrn-operon*) decreases. Finally, an increasing proportion of apparently unique, “novel” DNA, which shares no similarity to other Orobanchaceae genomes or any genomic sequences deposited in sequence repositories (Fig. 1A and Supplementary Table 3). Longer mitogenomes possess long stretches of unique DNA (e.g., 40 % in *Striga*), whereas the small- or middle-sized mitogenomes like those of *Castilleja*, *Lathraea*, *Rhinanthus*, and *Tozzia* contain less than 1%. To our surprise, the shortest mitogenome, *Melampyrum*, still have 12% of the total length identified as “novel DNA”. The majority of “novel DNA” remains unclear. We would like to point out that we cannot exclude that our limited representation of only 17 nuclear genomes across Orobanchaceae have led to underestimating the proportion of nuclear inserts. Hence, the proportion of true “novel” DNA might in fact be lower. Given our comprehensive sequence database search, we may conclude that those potentially missed true nuclear insert still represent Orobanchaceae or even lineage-specific unique DNAs.

### RNA editing in native and mosaic genes

Orobanchaceae mitogenomes have between 450 and 500 RNA editing sites in coding regions, all of them C-to-U conversions. Over 85% of the observed edits alter the amino acid due to changes in the first and second codon positions. RNA editing sites are unevenly distribution, with, for example, *atp1* showing no evidence for post-transcriptional nucleotide conversions and each of *ccmB*, *ccmC*, *mttB* and *nad4* containing more than 30 edit sites (Supplementary Table S6). In some species, RNA edits correct a mutated start codon in *nad4L* and *rps10* or restores stop codons as in *atp6*. However, it seems as if RNA editing in Orobanchaceae fails to rescue all mutations. For example, the *cox2* and *rpl10* in *Or. crenata* and *rps10* in *Ph. ramosa* all have extended reading frames due to substitutions in the stop codons that remain uncorrected.

We observe a slightly decreasing number of RNA editing site in a few obligate parasites, mostly holoparasites. Specifically, horizontally acquired genes and those showing footprints of conversion genes all have lost their conserved edit sites (e.g., *atp6* in *Phelipanche* and *Aphyllon*; *nad3* and *nad4L in Or. crenata*). Gains of editing sites or losses are otherwise rare, and if they occur, single nucleotide substitution appear to be causal. For example, T-to-C changes on the DNA level can create a new edit site, whereas substitutions of C to any other nucleotide results an edit site loss. We observe that such gains or losses occur several times independently across all investigated taxa and appear to follow no predictable order several times.

Mapping RNA data to the full mitogenomes showed that several noncoding regions, horizontally transferred DNA, as well as a few insertions from the nuclear genome are expressed at levels much lower than native mitogenes—especially in *Phelipanche* and *Striga*. Many of these expressed non-coding regions contain no notable or only a very short ORF. BLAST searches also failed to detect information of their potential function. It is noteworthy to mention that some noncoding regions, particularly in *Phelipanche* and *Striga*, are expressed at considerably higher coverages. Unlike the lowly expression regions, those RNA reads show several divergent bases to the mitogenome reference, including many G-to-A and C-to-U changes. We interpret these data not originating from mitochondrial genome, but from homologous regions in either the nuclear or plastid genome. This assumption is confirmed by further sequence analysis, which uncovered similarities to retrotransposons (e.g., LTR copia retrotransposon, identified by searching the Gypsy Database [Llorens et al. 2011]). The mitochondrial copy however is unlikely to be still active and expressed any more.

### Proliferation of repetitive mitogenomic DNA

Repetitive DNA constitutes up to 11%, is highly diverse in Orobanchaceae mitogenomes, and correlates significantly with overall mitogenome size (PGLS P-value < 0.001; Fig. 4A). Larger genomes (> 900 Kbp) like those of *Striga*, *Schwalbea*, and *Phelipanche* contain considerably more short repeats (SRs, 20–50 bp) than small mitochondrial genomes (PGSL P-value = 0.013). In contrast, repeat length in smaller mitogenomes depend mainly on large repeats of more than 100 bp (PGLS P-value = 0.021), most evident in *Conopholis*, *Lindenbergia*, *Melampyrum*, and *Triphysaria*. We classified repeats into three categories based on their length: short repeats, medium repeats (MRs, 50 – 100 bp) and long repeats (Supplementary Table S7). Together, the findings of opposite evolutionary trajectories of mitochondrial repetitive DNA imply that different molecular mechanisms underly the maintenance of small and large mitogenomes, whose size in turn relates to parasitic specialization (see above).

**Figure 4.**
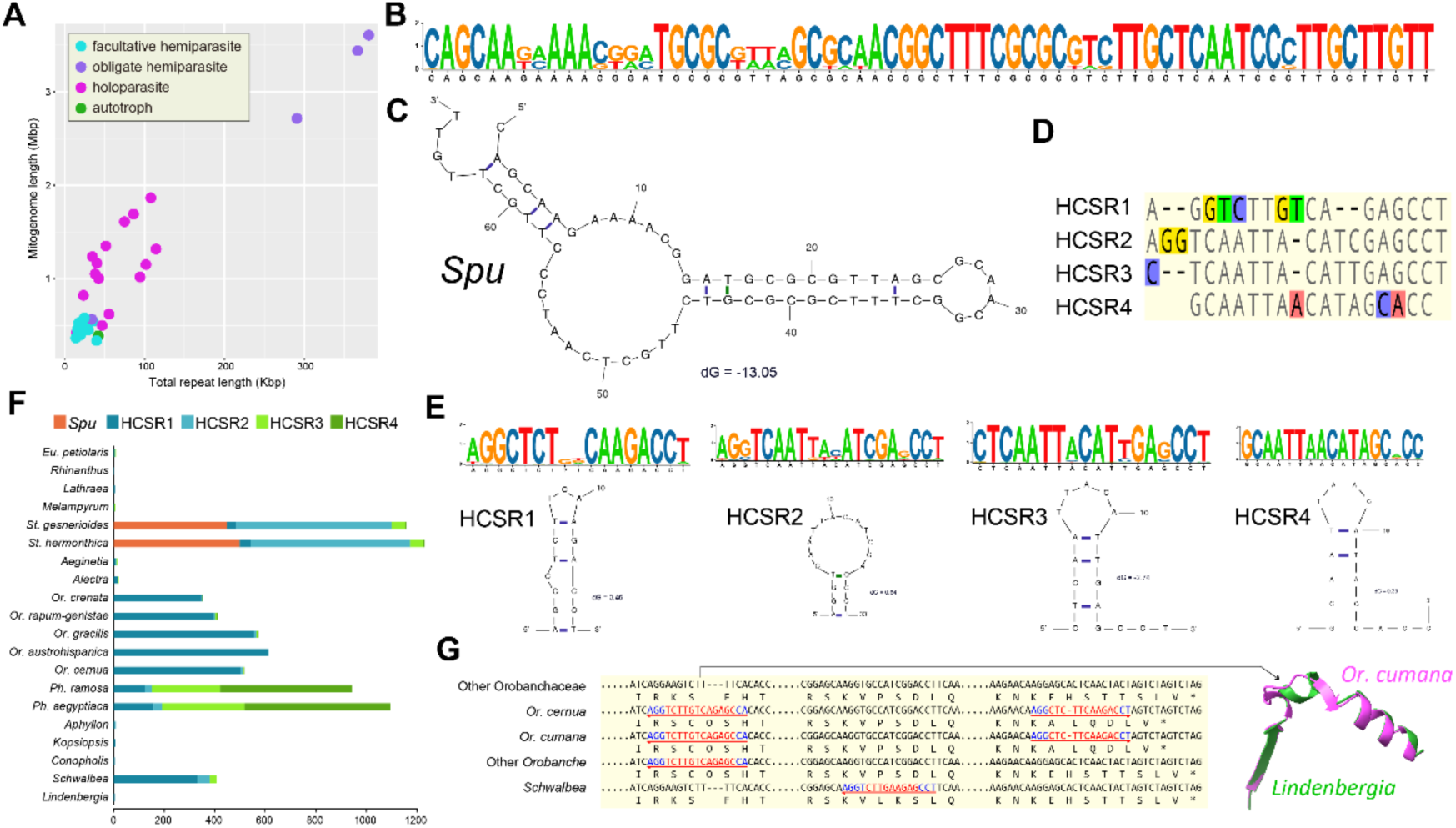
Repeat content and structural impacts of high-copy sequences in Orobanchaceae mitochondrial genomes. **(A)** Total repeat length is positively correlated with mitogenome size. **(B)** Sequence logo and **(C)** predicted secondary structure of the species-specific repeat *Spu*, based on all *Spu* sequences identified in *Striga* species. **(D)** High-copy sequence repeats (HCSRs 1–4) show substantial sequence similarity. **(E)** Sequence logos and predicted secondary structures for HCSRs. **(F)** Copy number of short repeats differs markedly among species. **(G)** Example of HCSR1 replacing intergenic regions within *matR*. Arrows beneath the sequences denote the extent and orientation of HCSR1; blue nucleotides indicate microhomology regions. Right panel illustrates predicted alterations in *matR* protein structure caused by a frameshift mutation in *Orobanche cumana* due to HCSR1 insertion.

**Figure 5.**
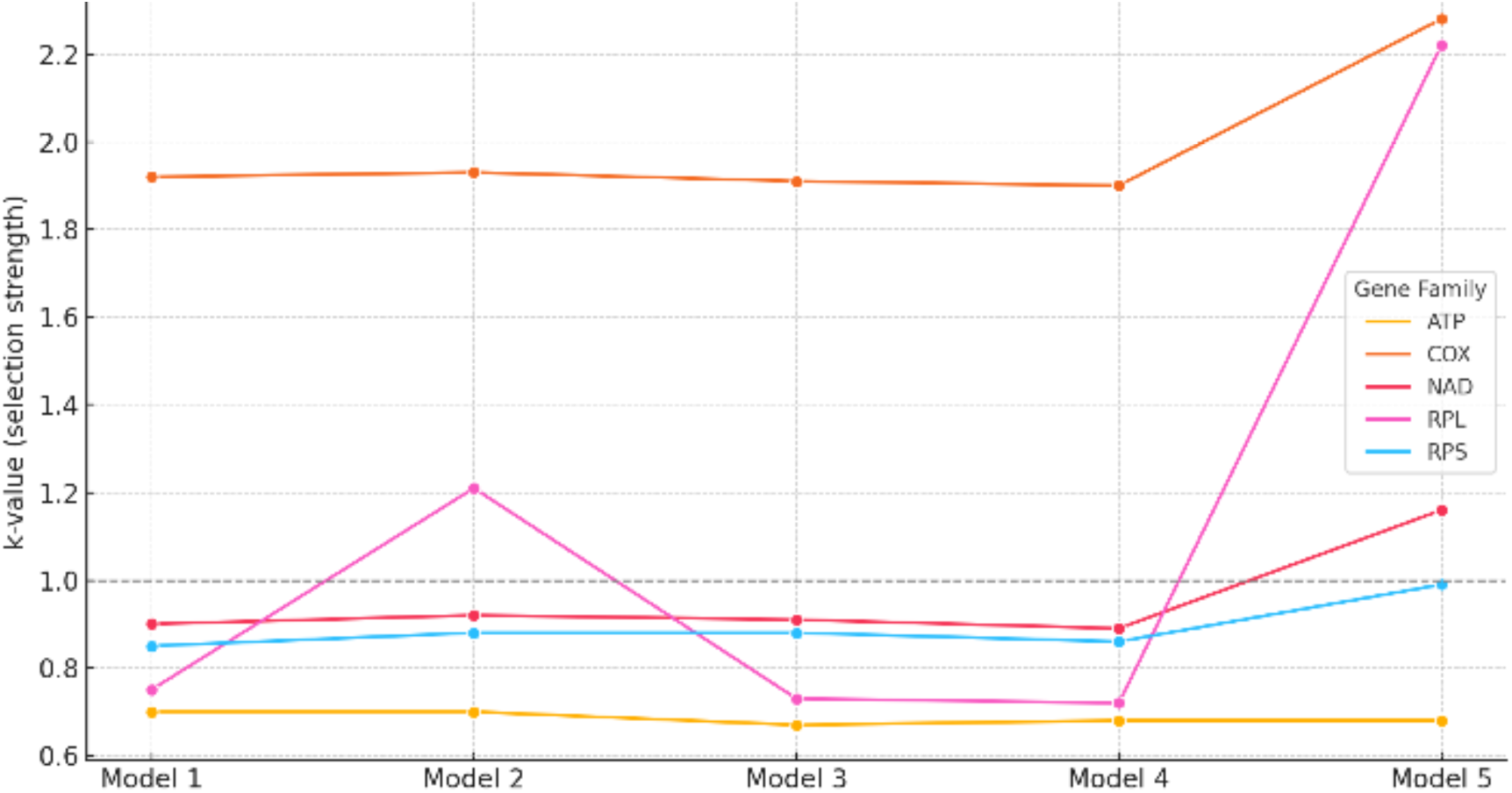
Distribution of selection strength (*k*) across mitochondrial gene classes in Orobanchaceae. Line plot showing variation in estimated selection intensity (*k*) across five evolutionary models for five key mitochondrial gene families: ATP synthase (ATP), cytochrome c oxidase (COX), NADH dehydrogenase (NAD), ribosomal large subunit proteins (RPL), and ribosomal small subunit proteins (RPS). Each model represents a different pairwise contrast of photosynthetic, non-photosynthetic, obligate, or facultative parasitic species within Orobanchaceae (Model 1: Photosynthetic vs. non-photosynthetic, with *Alectra & Striga gesnerioides* included in the latter); model 2: Facultative vs. holoparasitic (*Alectra* & *Striga gesnerioides* as non-photosynthetic); model 3: Cross-type switch 1 (*Alectra*: non-photosynthetic vs. *S. gesnerioides*: photosynthetic); model 4: Cross-type switch 2 (*Alectra*: photosynthetic vs. *S. gesnerioides*: non-photosynthetic); model 5: Obligate vs. non-obligate parasites). The dashed line at *k = 1* indicates neutral evolution; values <1 suggest relaxed purifying selection, while values >1 indicate intensified selection. COX genes consistently exhibit *k > 1*, indicative of positive selection across all contrasts, particularly in model 5. In contrast, ATP and RPS gene families generally display *k < 1*, consistent with relaxed selection in parasitic lineages. Model 5 also reveals intensified selection on NAD and RPL genes, suggesting shifts in mitochondrial functional constraint during the transition to obligate parasitism.

Small mitochondrial repeats are randomly dispersed, highly diverse in their sequence motifs, and present in high copy numbers—especially in Buchnereae (Supplementary Table S8). For example, the here named 66 bp-long core *Spu* repeat of *Striga* is replicated 447 and 499 times in *St. gesnerioides* and *St. hermonthica*, respectively (Fig. 4B). The different repeat copies’ identity is no less than 90% and subunits of it may occurs as palindromes (Fig. 4C). The *Spu* repeat and its variants resemble full-length or partial fragments from monocots, especially Poaceae—*Striga*’s preferred host plants (Supplementary Fig. S1). *Spu* inserts also occur in *rrnL* of *St. hermonthica*, but not in *St. gesnerioides*, and in *atp4*, *nad4*, and *rpl5* of both investigated *Striga* species, without causing an interruption of their transcript activity.

Apart from *Spu*, four smaller and sequence-related short repeats (HCSR1-4) of ∼20 bp are common in Orobanchaceae. In long mitogenomes, the copy number of these HCSRs can be in the hundreds, but their repeat motives also occur in smaller-sized mitochondrial genomes (Fig. 4D and Supplementary Table S8). Notably, they share a similar palindromic structure as *Spu* (Fig. 4E). These short repeats are also highly lineage-specific, showing different appetite in genera (Fig. 4F). Holoparasites *Aphyllon*, *Kopsiopsis*, and *Conopholis* were not found highly-copied short repeats. Interestingly, the short repeats could also insert into the coding regions and resulted in frame-shift mutations, which might further alter the protein structure (Fig. 4G). In sum, short repeats, that in part originate from host plant mitochondrial DNA shows an extraordinary dynamic in Orobanchaceae genomes and contribute to mitochondrial size inflation.

### Selection pressure on mitochondrial genes in parasitic and non-parasitic Orobanchaceae

To investigate selection pressures acting on mitochondrial genes in parasitic plants, we applied the RELAX framework (Wertheim et al. 2015) to five comparative models (Figure 5; see below – *Materials and Methods*). The selection intensity parameter (*k*) was used to infer relaxed (*k* < 1) or intensified (*k* > 1) selection, with statistical significance assessed through p-values.

Several ATP synthase genes exhibited significant evidence of relaxed selection across all tested models (Supplementary Table S9). The concatenated ATP dataset showed consistently low *k* values (∼0.7, p < 0.001) across all comparisons, indicating widespread relaxation of selection pressure. Individually, *atp1* displayed significant relaxation (*k* ∼0.6, p < 0.05) in all models, while *atp8* showed nearly significant relaxation (*k* = 0.61, p = 0.0596) in obligate parasites. These findings suggest a functional reduction of ATP synthesis in parasitic species, perhaps due to a reliance on host-derived energy sources.

Ribosomal genes also exhibited clear evidence of relaxed selection, particularly within the large ribosomal subunit (RPL). The concatenated RPL dataset (*k* = 0.72–0.75, p < 0.05) and *rpl2* (*k* = 0.33–0.39, p < 0.001) showed strong relaxation across multiple comparisons. Additionally, *rpl10* (*k* = 0, p < 0.1) suggests a potential loss of function. The RPS dataset also showed moderate relaxation, with RPS (p = 0.0264– 0.0433) displaying significantly reduced selection pressure. The pattern of relaxed selection across ribosomal genes aligns with prior findings indicating a reduction in mitochondrial ribosomal functionality in parasitic plants.

Conversely, genes associated with core mitochondrial functions, such as cytochrome c oxidase (COX) and NADH dehydrogenase (NAD), exhibited intensified selection. The concatenated COX dataset displayed consistently elevated *k* values (∼1.9, p < 0.001) across all comparisons, indicating strong selective pressure. Among individual COX genes, *cox1* showed an unexpected signal of relaxation in obligate parasites (*k* = 0.62, p = 0.0194), suggesting lineage-specific variation in selective constraints. Among NADH dehydrogenase genes, *nad6* displayed significantly intensified selection in obligate parasites (*k* = 2.75, p = 0.0437), while *nad5* (*k* = 1.35, p = 0.1652) showed a similar trend but lacked statistical significance. The observed pattern suggests that certain NADH dehydrogenase components remain under strong selection, potentially reflecting adaptations to altered electron transport chain function in parasitic species. Additionally, genes associated with cytochrome c maturation showed evidence of selection intensification. *ccmFn* exhibited a highly significant increase in selection intensity (*k* = 1.57, p = 0.0002) in obligate parasites, highlighting its potential importance in mitochondrial function despite overall genomic streamlining.

The observed selection patterns align with the hypothesis of mitochondrial genome streamlining in parasitic plants, where genes related to ATP synthesis and ribosomal function are experiencing relaxed selection, while genes essential for electron transport and core mitochondrial processes remain under selection pressure. The complete loss or extreme relaxation of *rps14* (*k* = 0, p < 0.02 across models) and *rpl10* (*k* = 0, p < 0.1) suggests pseudogenization or functional loss of these ribosomal components. Furthermore, while most ribosomal and ATP synthase genes showed relaxed selection, exceptions such as *matR* (k = 1.44, p = 0.001) in obligate parasites indicate selective retention of specific mitochondrial functions. In sum, these findings provide strong evidence for divergent selection pressures in parasitic plant mitochondrial genomes, with relaxed selection acting on ATP synthase and ribosomal genes, while core mitochondrial functions, including cytochrome c oxidase and NADH dehydrogenase, remain under intensified selection. These patterns are consistent with a transition toward host-dependent energy metabolism and mitochondrial genome streamlining in parasitic species.

## Discussion

Our analysis of mitochondrial, plastid, and nuclear genome evolution across 30 species of Orobanchaceae revealed a strong coevolution of all genomic compartments along the transition from a photosynthetic to a nonphotosynthetic lifestyle. Similar trends of mitogenome size inflation in holoparasitic plants compared with closely related nonparasitic species occur in *Cynomorium* (Cynomoriaceae; Bellot et al., 2016), *Lophophytum* (Balanophoraceae; Sanchez-Puerta et al., 2017), *Cuscuta japonica* (Convolvulaceae; Lin et al., 2022), and *Gastrodia* (Orchidaceae; Yuan et al., 2018). In contrast hemiparasites and physiological holoparasites like *Viscum scurruloideum*, *Rhopalocnemis phalloides*, or other *Cuscuta* possess small or medium sized-mitogenomes–resembling the evolutionary patterns observed in Orobanchaceae (Skippington et al. 2015; Lin et al. 2022; Yu et al. 2022). It remains to be investigated whether these parasite lineages also show a coevolution between organellar and the nuclear genome with regard to structural and functional evolution must be studied in more detail.

As plastid genomes shrink in size (and functionality), both the nuclear and mitochondrial genome gain length. Mitogenome evolution exhibits obvious lineage-specific features regarding their architecture, coding capacity, and complexity of the gene and DNA mosaic (Fig. 1A). With this being the first systematic study covering all intracellular genomes in parasitic plants, existing data does not allow presenting a conceptual model of the mechanisms underlying the coevolution of genomic evolution along lifestyle transitions. Based on reports that targeted single genomic compartments (reviewed in Wicke and Naumann 2018), we may conclude that the transition to parasitism and the functional reconfiguration of the plants genetic repertoire triggers or proceeds as a feedback loop across all genomic compartments. For example, relaxation of selectional pressures on plastid genomes in parasites, might also affect the selectional regimes of proteins maintaining the plastid genetic systems, resulting in a slight relaxation of their selection pressure. In consequence, nuclear-encoded repair and other housekeeping proteins targeted to both organelles, of which there are many, could lose efficiency and cause more dramatic sequence and genomic change in parasites. Our data of minimal changes in physiological holoparasites in comparison with their closest photosynthetic relative in Orobanchaceae suggest a time lag before functional reductions of plastid genomes take effect in other genomic compartments. A denser taxon representation of taxa of distinct parasitic specialization and genome-wide ancestral genome reconstruction based on chromosome-level sequencing might uncover the extent, directions, and evolutionary tempo of lifestyle-genotype associated genomic change and their underlying mechanisms.

Contributing to mitogenome inflation are increasing proportions of integrated intracellularly transferred DNA and genetic material from other (host) plants. Besides, repeat proliferation is associated with mitogenomic growth as well as an expansion of unique and apparently “novel” DNA, whose origin remain unknown or cannot be determined precisely due to limitation regarding available genomic data. All size-influencing factors are not only correlated with genome size, but also with the investigated plants’ lifestyles, revealing their maxima in obligate parasites. Although nuclear genomic fragment become more abundant in holoparasites and the derived obligate parasites of *Striga*, plastid insertions are more abundant and longer in (many) facultative hemiparasites. Known from *Lophophytum* (Sanchez-Puerta et al., 2017) as well, this puzzling finding, may be explained by more recent and frequent incorporations of raptured plastids in photosynthetic parasites, which are also thought to contain more plastids or plastome copies in a cell (Wicke et al. 2013; Feng et al. 2016).

The more intimate the contact within parasite-host systems become, as it happens along parasitic specialization in plants, the more exposure to the genetic material of hosts is a given for the parasite. Therefore, it is not surprising that horizontal gene transfer from hosts is a well-known feature of parasitic plant genomes (Davis and Wurdack 2004; Mower et al. 2004, 2010; Barkman et al. 2007; Yang et al. 2016, 2019; Sanchez-Puerta et al. 2017, 2019; Skippington et al. 2017). Here, we show the extent of foreign DNA integration and how it correlates with mitogenome expansion in parasitic Orobanchaceae. Mitogenome-wide integration of genetic material from a host plant was reported for the holoparasite *Lophophytum* (Balanophoraceae) as well (Sanchez-Puerta et al. 2019). The long fragmental coverage of host-like DNA corroborated mitochondrial fusion as involved mechanism. In Orobanchaceae, shorter fragments of foreign genetic material dominate the mitogenomic landscape. We may assume that direct uptake of non-native DNA might be another important mechanism. However, integration following mitochondrial fusion with subsequent fragmentation of newly inserted DNA is likely, too. Despite normally low nucleotide substitution rate in plant mitogenomes (Wolfe et al. 1987; Palmer and Herbon 1988), streamlining incorporated DNA to match mitogenomic nucleotide composition may be important for genomic maintenance and a prevalence of posttranscriptional C-to-U edits (Table S3,). Understanding this process in detail might allow using the insertions as internal genomic clocks to estimate fragmental survival and turnover times in mitochondrial DNA.

Rapid and dynamic evolution of the overall genomic architecture of parasitic plant mitogenomes involves its coding fraction (Supplementary Table S2). Orobanchaceae, and other parasitic plants, utilize immigrant or chimeric genes of full-length or partial host origin, and re-gain formerly lost genetic material by complementing their own coding capacity with that of their hosts (Mower et al. 2004, 2010; Xi et al. 2012; Bellot et al. 2016; Sanchez-Puerta et al. 2019). Associated with host-derived gene fragments, otherwise conserved RNA editing site losses occur in Orobanchaceae. Our data of the gene incorporation history suggest that those edit site losses are rather due to a DNA-based replacement of native through host gene fragments and/or gene conversion rather than retroprocessing. Unfortunately, the complex history of the parasites’ mitogenomes and its coding fraction hampers substitution rate inferences. Nonetheless, future studies centering on patterns of sequence drift during mitogenomic evolution along parasitic specialization will be worthwhile and might uncover strongly divergent patterns of substitution rates and selectional footprints than known from plastid DNAs of parasitic plants (Wicke et al. 2016; Wicke and Naumann 2018).

Mechanistically, incorporation of foreign DNAs into mitochondrial genomes of plants may be facilitated by the organelle‘s active import of DNA (Konstantinov et al. 2016), paired with organelle fusions and fissions (Arimura et al. 2004; Sheahan et al. 2005). Experimental evidence has shown that DNA translocation of favorably shorter, linear fragments across the mitochondrial membrane is unspecific and independent of the original sequence (Koulintchenko et al. 2003; Ibrahim et al. 2011). Hence, we may conclude that the incorporation of DNA fragments from other intracellular genomes of other species swimming in the cytosol is equally likely. Besides, gains of genetic material, functional or not, can occur via mitochondrion-to-mitochondrion fusions between hosts and parasites (Rice et al. 2013; Sanchez-Puerta et al. 2017), the latter of which can also result in parasites containing horizontally acquired DNA from non-host plants (Gandini and Sanchez-Puerta 2017), which we also observe here. Alternatively, the low substitution rate of plant mitogenomes could freeze foreign DNAs for a very long time, and, thus, potentially reflect host switch histories.

Changes of the mitochondrial repeat diversity (Supplementary Table S7) over the gradual physical growth of mitochondrial genomes towards holoparasitism is another important contributor to genomic change. In itself, repeat proliferation already provides further clues of the mechanism underlying the changes of the parasites’, and other plants’ genomic landscape. Especially short repeats can have a strong influence on genome architecture and stoichiometry by mediating recombination or rearrangement processes (André et al., 1992; Kanazawa et al., 1998; Allen et al., 2007; Shedge et al., 2007; Alverson et al., 2011)(André et al. 1992; Kanazawa et al. 1998; Allen et al. 2007; Shedge et al. 2007; Alverson et al. 2011). To maintain genomic integrity, plastid DNA normally selects against repeat proliferation (Wicke et al. 2013)—unlike mitogenomes that might even use repeats to generate structural dynamics and genetic novelty. The abundant *Spu* elements of monocot origin in *Striga* mirrors patterns of short-repeat proliferation reported in extremely large angiosperm mitogenomes, although not all large mitogenome are rich in short repeats (Gandini et al. 2019). Particular *Spu* and other repeat motifs with a hairpin palindromic structure (Fig. 4) appear to constitute a mobile element of some seed plant mitogenomes (Nakazono et al. 1994, 1995; Paquin et al. 2000; Aono et al. 2002; Chaw et al. 2008; Erpenbeck et al. 2009; Lavrov 2010), perhaps by partial reversed transcription of mitochondrial transcripts (Nakazono et al., 1995; André et al., 1992; Gandini et al., 2019)(André et al. 1992; Nakazono et al. 1995; Gandini et al. 2019). In Orobanchaceae, short repeats locate near foreign DNA insertions, hinting at a potential association with those structural changes irrespective of how the genetic material has entered the organelle. Short repeats facilitate DNA breaks, which not only provides the opportunity to integrate foreign DNAs, but create recombination sites and trigger their own proliferation through erroneous DNA repair. Not surprisingly, thus, repeat landscape reshaping associates with MMEJ that mediates the repair of DNA strand breaks via short homologous bases (< 10 bp) at the expense of short deletions (McVey and Lee 2008; Chiruvella et al. 2013), which we commonly observed in the investigated taxa (e.g., Fig. 3C and 4G). In sum, recombination as well as proliferation of short-repeats paired with microhomology-based repair appears to represent dominating mechanisms of the observed intra- and intergenic variations in Orobanchaceae, likely acting also in nonparasitic plants.

## Materials & Methods

### Taxon sampling and sequencing

We sequenced, assembled, and analyzed the complete mitochondrial and plastid genomes of 30 Orobanchaceae, including an autotrophic representative (*Lindenbergia philippensis*), 12 hemiparasites (*Euphrasia minima*, *E. petiolaris*, *Melampyrum pratense*, *Neobartsia pedicularoides*, *Odontites vernus*, *Phtheirospermum japonicum*, *Rhinanthus serotinus*, *Schwalbea americana*, *Striga hermonthica, Tozzia alpina*, *Triphysaria versicolor,* plus *Castilleja paramensis* that was presented earlier by Fan et al., 2016), and 15 holoparasites (*Aeginetia indica*, *Aphyllon californicum*, *Conopholis americana*, *Kopsiopsis hookeri*, *Lathraea clandestina*, *Orobanche austrohispanica*, *Or. cernua*, *Or. crenata*, *Or. cumana*, *Or. densiflora*, *Or. gracilis*, *Or. minor*, *Or. rapum-genistae*, *Phelipanche aegyptiaca* and *Phe. ramosa*). The two physiological holoparasites *Alectra orobanchoides* (De La Harpe et al. 1979, 1981) and *Striga gesnerioides* (Graves et al. 1992) complemented our sampling to narrow down the relative evolutionary timing of lifestyle-depending genomic changes. DNA was extracted from fresh or silica-dried material using previously described methods for Orobanchaceae (Wicke et al. 2013, 2016) and subjected to whole-genome shotgun sequencing on an Illumina HiSeq2000 or HiSeq2500 in paired-end mode (Supplementary Table S10).

### Sequence data processing and genome assembly

All shotgun data were quality-trimmed with *Trimmomatic v0.36* (Bolger et al. 2014), admitting only sequences of at least 70 bp with a minimum Phred score of 30 to enter downstream analyses; overlapping read pairs were merged and identical reads removed. All nuclear drafts were assembled using SPAdes v3.10 (Nurk et al. 2013) with k-mers 21, 33 and 55 and *CLC Assembly Cell* (kmer 33; bubble size: 50; contigs updated after read back-mapping); contigs smaller than 200 bp were removed). Quality of the different assemblies was assessed using Quast v.4.5 (Gurevich et al. 2013) under default parameters for eukaryote genomes, and the higher-quality assemblies per species were selected for further analysis.

For the target assembly of organelle genomes, we randomly extracted between two and six Gbp from each of the cleaned raw datasets and assembled these subsets in VELVET v1.2.10 (Zerbino and Birney 2008) with five kmer ranges from 57 to 87, automatically adjusting the expected coverage as well as the coverage cutoff; contigs longer than 200 bp were saved. Plastomes were assembled exactly as described earlier (Feng et al. 2016). The VELVET contig pools per species were sorted by their similarities to 106 mitogenomes of angiosperms using *megablast* v2.40+ at default parameters, and contigs with hits of an e-value better than 1e-10 were extracted. The plastome of *Lindenbergia philippensis* was used to mask plastid contigs. Identified mitogenome candidates then were connected manually, for which 70 to 90 bp of the contig ends were searched against the contig pool and the species-specific plastome. Contigs were connected directly when their terminals matched. If no matching end was retrieved in the mitochondrial or plastid-like contigs, we expanded the search to the full contig pool, and best-matching ends were connected manually. When contig ends matched with sequences residing inside of a contig, we considered these as potential repeats. To resolve those repeats, we mapped reads iteratively to candidate contigs, and determined the repeat boundaries by coverage, screened for diverging reads at the boundaries and used these as extension primers. If contig ends matched a plastome regions, we marked its position and direction and proceeded similar to resolving repeats albeit with different coverage cutoffs. These steps were iterated until no further contig extension was possible or a circular molecule topology was obtained. Finally, paired reads were mapped back to the genomes to check and correct potential misassemblies based on coverage, GC content, and conflicting read orientation information of sequence pairs.

### Annotation

We used the published mitogenomes of *Castilleja paramensis* (GenBank ID: NC_031806), *Mimulus guttata* (GenBank ID: NC_018041) and, then incomplete, *Neobartsia pedicularoides* (GenBank ID: KP940485–KP940493) as references to annotate coding and rRNA genes manually by similarity in *GENEIOUS* R10 (Biomatters, Inc.,) in addition to using *Mitofy* (Alverson et al. 2010). tRNAscan-SE v1.3.1 (Lowe and Eddy 1997), as implemented in *Mitofy*, was employed to detect and annotate tRNAs.

Coding genes with disrupted reading frames, premature stop codons, or non-triplet frameshifts were annotated as pseudogenes. A graphical summary of the gene content was generated using *BRIG* v0.95 (Alikhan et al. 2011) with *Lindenbergia*’s mitochondrial gene set as the reference. The protein structures were predicted by AlphaFold 3 server (Abramson et al. 2024) and the visualization and comparison were done in UCSF ChimeraX v1.7.1 (Meng et al. 2023).

### Nuclear genome size estimates

Unless data existed from previous studies (Weiss-Schneeweiss et al., 2006; Piednoël et al., 2012; Wicke, 2013)(Weiss-Schneeweiss et al. 2006; Piednoël et al. 2012; Wicke 2013), we estimated the genome sizes of our study taxa by kmers. After trimming all bases with a quality of 20 or lower, we employed *Jellyfish* v2.2.4 (Marcais and Kingsford 2011) to count 21-mers. The kmer-distributions then were graphically summarized and the genome size was estimated as total number of k-mers divided by the peak position of the kmer distribution. Where available, we compared estimates obtained with different methods to estimate the error of kmer count-based genome size estimation using low-coverage assemblies; we observed only minor divergences.

### Dispersed repeats

Forward and reverse compliment repeats were detected by using blastN (word size: 11, default gap open and extension penalties). Repeat length and number were counted by using a custom Perl script based on three size categories, 20 – 50 bp, 50 – 100 bp and longer than 100 bp. Only identity greater than 95% were considered. Overlapping bases of different repeat motifs were summed only once in length. The highly copied short repeats were found by an additional assembly of the repeats then searched manually. Their secondary structures were predicted by Mfold web server (Zuker 2003).

### Analysis of mitogenomic mosaics

We blastN-searched the mitogenomes of each species against the mitochondrial DNAs (mitoDNA) of the other Orobanchaceae study taxa. Hits with an e-value of at least 1E-10 were considered significant. Only regions of at least 200 bp in length that had hits in at least half of the included genera (instead of species to avoid the bias of single or multiple sequenced genera) (i.e., min. 10 genera in Orobanchaceae) were classified as “conserved homologous mitochondrial DNA” (CH). We considered all hits from at least two (including the same) and less than half of all included genera as “nonconserved” (NCH). Plastid-derived, HGT, Nuclear-derived DNAs were masked (see below) from both CH and NCH. Subsequently, we blastN-searched the mitogenomes of our study taxa against 106 mitogenomes, 1444 plastomes, 39 nuclear genomes of flowering plants, and whole-genome shotgun assemblies of our Orobanchaceae species. Custom Perl scripts were used to rank the results, sorting by length, similarity, and e-value for a detailed inspection (see below). Regions of at least 200 bp with no hits to any of the reference genome were preliminarily flagged as “novel DNA”, which we re-analyzed against the full, non-redundant NCBI nucleotide database (21^th^ March, 2018) and processed the results as above.

To analyze specifically plastid-derived fragments, we considered regions with high similarity (min. evalue: 1e-10) to plastid DNA with a length of 100 bp or more as plastid-derived DNA (MTPU). Of these, we classified fragments with significant hits to Lamiales species as ‘intracellular DNA transfer’, whereas those with similarity to distantly related taxa were treated as candidates of horizontally acquired DNA. We ignored hits to the mitochondrial-derived plastid DNAs of two actual host species (Iorizzo et al. 2012; Straub et al. 2013).

Mito-derived fragments were evaluated using a blastN-based assessment of foreign mitoDNA. We limited our analysis to noncoding sequences between mitochondrial coding genes and/or plastid-derived fragments; coding regions (included duplicated genes) where analyzed as described below. We considered only continuous best-hit regions of more than 200 bp in length as significant, ranked the results by similarity, hit length, and taxonomic affiliation, whereby hits by family corresponding to less than 70% of the matching fragment were discarded. *Lindenbergia* was treated as an outgroup to the parasitic Orobanchaceae. To minimize false-positive hits regarding the identification if HGT donors, hit regions of parasitic Orobanchaceae were soft-masked temporarily. For relevant matches to more than three families, we built phylogenetic trees using maximum likelihood inferences as described below. If Orobanchaceae sequences clustered with other Lamiales species, irrespective of potential gene tree/species incongruency, we interpreted this as evidence against HGT. Parasite fragments placed within non-Lamiales clades were considered as potential HGT event, for which we determined the donor as the nearest-affiliated family of said fragment. The confidence level of these individual mitogenome-to-mitogenome HGT events was ranked as low, medium, or high by comparing the match or relatedness of the retrieved donor fragment to the known host species of the investigated parasite species and genera; the total length of each mito/mito HGT confidence categories per species was summed up. Matches to *Lindenbergia* or Phrymaceae were considered as due to shared ancestry. Lentibulariaceae, Gesneriaceae, Lamiaceae, and Oleaceae are closely related to Orobanchaceae but some species of those families are known also as hosts of some herein investigated parasites, and only in these cases, their lengths were included in ‘host length. Direct matches of HGT candidates to known host lineages were classified as “high-confidence”, while those to non-host taxa as “medium confidence”. We performed a case-by-case examination for all parasites for which no host lineage was represented in the reference database (e.g., for *Conopholis americana*, because no Fagaceae mitogenome was publically available as of writing this article). We classified hits from Amborellaceae, Balanophoraceae, Cynomoriaceae, and Santalaceae as “unclear” origin, because representatives of either of these families are known to contain genes acquired horizontally themselves (Rice et al. 2013; Bellot et al. 2016; Sanchez-Puerta et al. 2017; Skippington et al. 2017). The plot was generated in R with ape (Paradis et al., 2004) and phytools (Revell 2012) packages.

Nuclear genome-derived fragments were analyzed based on 16 shotgun-sequenced genome drafts of our Orobanchaceae species. The remaining 14 species were not considered here because their average nuclear genome coverages were too low (< 7X). We removed organellar sequence reads based on matches of the organelle genomes and their coverages. To further investigate the interplays of mito- and nuclear genomes as well as the potential nuclear HGT level, we combined the other 23 published nuclear genomes representing the major host lineages of our study parasites and mitogenomes of lamiids. Identifying regions of nuclear genomic origin essentially followed the same routines and used the same criteria for classification as described above, with the exception that all mitochondrial HGT candidates were masked from the queried noncoding sequences. To account for genome quality differences and the considerably smaller reference taxon set, we considered also hits from other Orobanchaceae, Phrymaceae, and Pedaliaceae as potentially originating from the parasites’ own nuclear genome. Regions containing significant hits (>100 bp, evalue < 1e-10) only from genome(s) other than the above-mentioned families were considered as HGT candidates (with medium confidence).

### Analysis of compositional biases

To inspect changes of nucleotide composition (GC content) following intracellular gene transfer plastid-like fragments, we blast-searched the mitogenome of each species against its own and the *Lindenbergia* plastome. Since the plastid nucleotide composition also changes along reductive plastome evolution towards holoparasitism (meta-analyzed by Wicke and Naumann 2018), we can approximate from GC content comparisons of the own vs. an “ancestral” autotrophic plastome, when—in an evolutionary context—the transfer might have occurred. We limited our analysis of fragments of at least 100 bp and inferred the GC content of mitogenomes’ plastid-derived fragments and their corresponding plastid sequences with help of a custom Perl script. We analyzed the direction and significance of compositional changes by generalized linear mixed models, where species were treated as random factor.

### ORF searching and functional prediction

Open reading frames (ORFs) were extracted when their lengths were or exceeded 50 aa under standard genetic code constraints and the start codons ATG, TTG or CTG where detected. All identified ORFs were passed on to *InterProScan* v. 5.29-68.0 (Finn et al. 2017) to predict potential proteins with default parameters. Result were categorized into six classes: mobile element-like (including reverse transcriptase-like domains, integrases, transposases, RNAse H, and proteins associated with terms such as gag-pol, viral or retroviral, retropepsin, retrotransposon; hits to “mitovirus” was counted separately), domain of unknown function, zinc domains (incl. zinc finger, CCHC-type, zinc knuckle), prokaryotic-like (incl. pilin assembly, prokaryotic membrane lipoprotein lipid attachment site profile, TraU, mating pair stabilization), phage-like proteins (incl. bacteriophage head to tail connecting protein, tail tubular protein, baseplate J-like protein, phage Mu protein F like protein, phage lysozyme, phage regulatory protein, phage terminase, phage term 2, PBSX family, DNA-packaging protein), and others.

### Analysis of mitogenome expression

We selectively analyzed the expression of mitochondrial genes to estimate the extent of RNA editing and validate mitochondrial gene models. In addition to readily available RNAseq data for *Aphyllon californicum*, *Phelipanche aegyptiaca*, *Striga hermonthica*, and *Triphysaria versicolor* (Wickett et al. 2011; Yang et al. 2015, 2016), we isolated total RNA of *Aeginetia indica, Lindenbergia philippensis, Rhinanthus serotinus, Lathraea clandestina, Orobanche crenata, Phe. ramosa*, *Phtheirospermum japonicum,* and *Striga gesnerioides* using the Ambion™ PureLink® RNA Mini Kit according to the manufacturer’s instructions; two isolations per species were prepared. Sequencing was performed on an *Illumina* HiSeq2500 in 150 bp paired-end mode. RNAseq data were subjected to quality trimming using *Trimmomatic* as above, before we mapped the cleaned data first to the species-specific similarity-based predictions of coding sequences per protein-coding gene and, secondly, to the entire mitogenome per species in GENEIOUS. RNA editing sites were identified by pairwise alignment of the coding sequence (DNA) to the consensus sequences of the RNAseq mappings, whereby the consensus was built by 25 % majority rule. If their average coverage exceeded 5X, transcribed non-coding or previously unannotated regions were passed on to exploratory blastN and blastX searches for further characterization and annotation.

### Phylogenetic analyses

For species tree reconciliation, every organellar protein-coding gene was first codon-aligned with *PAL2NAL* (Suyama et al. 2006) under default settings, then optimized with *Guidance*II (Sela et al. 2015) and the *MAFFT* aligner v7.306b (Katoh and Standley 2013) in codon-mode, and finally concatenated to an organelle gene dataset. Using *Hyphy* v2 (Kosakovsky Pond et al. 2005), we optimized an unconstrained likelihood function with local parameters for a GTR-Gamma model, a guide tree representing the generally established species relationships of our study taxa according to Schneeweiss (2013), and the concatenated organelle gene data to obtain its maximum likelihood. The resulting phylogenetic tree was exported with branch-length information as basis for phylo-statistical analyses.

To gather further evidence for HGT, mitochondrial genes and all HGT candidates, irrespective of their alleged genomic origin, were analyzed separately, for which all fragments were sorted into homologous nucleotide data sets; sequences departing by more than 30 % from median length were discarded. All data sets were aligned with *MAFFT* v7.306, and phylogenetic trees were built from each of the resulting aligned dataset using RAxML v8.2.4 (Stamatakis 2014) with the GTR-GAMMA model. We used *R* and the *ape* package (Paradis et al. 2004) for graphical illustration of the resulting gene trees.

## Data availability

The newly assembled organelle genomes have been submitted to NCBI Genbank and will be released prior to publication. Additional datasets related to our HGT, MTNU, and InterProScan analyses are deposited on FigShare: https://figshare.com/s/98817f4e000276ab2593.

## Author contributions

S.W. conceived the project. S.W. and Y.F. together performed the experiments, analyzed the data and wrote the manuscript.

