## Supplementary figures and images for "Systemic organellar genome reconfiguration along the parasitic continuum in the Broomrape family (Orobanchaceae)"

### SupplementaryFig_S1.pdf

Sequence Logo

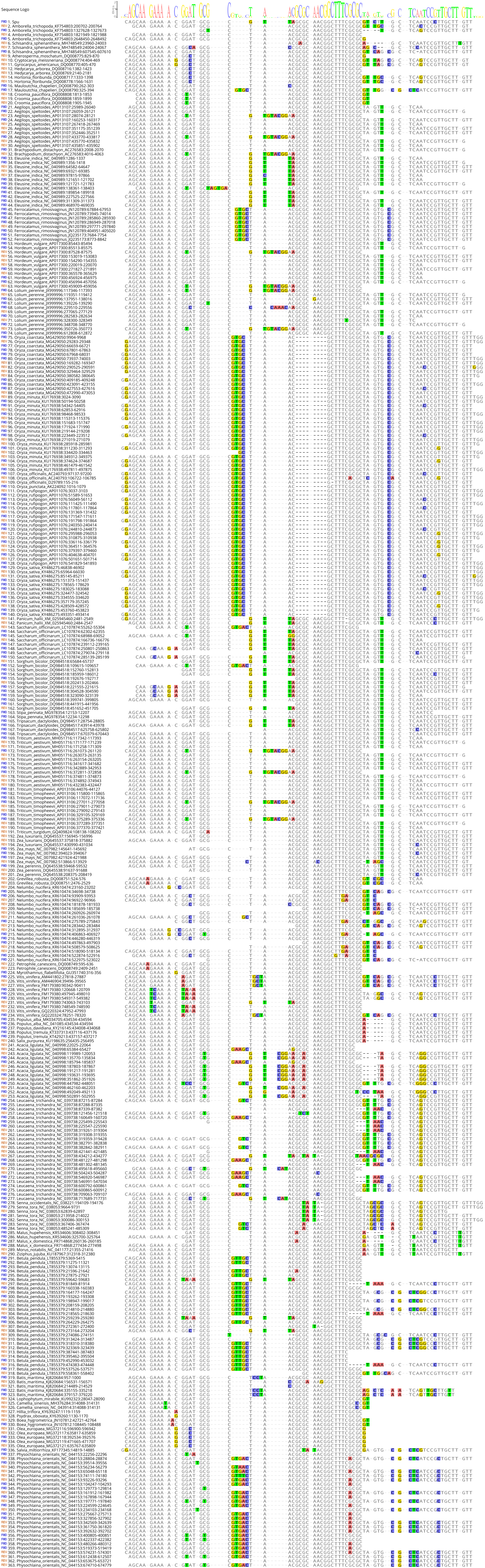
